# A Synthesis of Game Theory and Quantitative Genetic Models of Social Evolution

**DOI:** 10.1101/2021.03.27.437341

**Authors:** Joel W. McGlothlin, Erol Akçay, Edmund D. Brodie, Allen J. Moore, Jeremy Van Cleve

## Abstract

Two popular approaches for modeling social evolution, evolutionary game theory and quantitative genetics, ask complementary questions but are rarely integrated. Game theory focuses on evolutionary outcomes, with models solving for evolutionarily stable equilibria, whereas quantitative genetics provides insight into evolutionary processes, with models predicting short-term responses to selection. Here we draw parallels between evolutionary game theory and interacting phenotypes theory, which is a quantitative genetic framework for understanding social evolution. First, we show how any evolutionary game may be translated into two quantitative genetic selection gradients, nonsocial and social selection, which may be used to predict evolutionary change from a single round of the game. We show that synergistic fitness effects may alter predicted selection gradients, causing changes in magnitude and sign as the population mean evolves. Second, we show how evolutionary games involving plastic behavioral responses to partners can be modeled using indirect genetic effects, which describe how trait expression changes in response to genes in the social environment. We demonstrate that repeated social interactions in models of reciprocity generate indirect effects and conversely, that estimates of parameters from indirect genetic effect models may be used to predict the evolution of reciprocity. We argue that a pluralistic view incorporating both theoretical approaches will benefit empiricists and theorists studying social evolution. We advocate the measurement of social selection and indirect genetic effects in natural populations to test the predictions from game theory, and in turn, the use of game theory models to aid in the interpretation of quantitative genetic estimates.

## Introduction

Social interactions are ubiquitous in nature (Frank 2007). In nearly every species, conspecifics interact in some way during their life cycle in contexts such as territoriality, mating, and parental care, and these social interactions can have profound implications for evolution (West-Eberhard 1979, 1983; Moore et al. 1997; Frank 1998). Darwin (1859) realized the importance of social context for evolution early on, and proposed the mechanisms of sexual selection and family-level selection to account for elaborate courtship displays and non-reproductive castes in social insects, respectively. However, when theoretical population genetics began to develop in the twentieth century to provide a mathematical basis for Darwin’s theory, it was slow to incorporate social evolution. Stalwarts of the Modern Synthesis like Fisher (1930) and Haldane (1932) described some scenarios involving social evolution qualitatively but did not offer a mathematical analysis (Karlin and Matessi 1983).

In the 1960’s, behavior began to be viewed separately given its unique property as a both a target and agent of selection, attracting the attention of formal mathematical theory. Hamilton’s (1963; 1964a; 1964b) seminal work, which showed how genetic relatedness and social fitness effects combine to predict the evolution of social traits, catalyzed the development of the mathematical theory of social evolution and the fields of behavioral ecology and sociobiology in the decades to follow. Hamilton’s early models of the evolution of altruistic behavior were explicitly genetic, and his concept of inclusive fitness involved viewing selection through what was later termed the “gene’s eye view” (Dawkins 1976). However, later developments in social evolution theory often ignored genetics and relied on the “phenotypic gambit” (Grafen 1984), the assumption that the approach to phenotypic evolutionary optima is usually unconstrained by genetics (Eshel 1996). The most influential theoretical approach to phenotypic social evolution was evolutionary game theory and its central concept, the evolutionarily stable strategy (ESS) (Maynard Smith and Price 1973; Maynard Smith 1982; Dugatkin and Reeve 1998; Hofbauer and Sigmund 1998; McElreath and Boyd 2007; McNamara and Leimar 2020). Evolutionary game theory offered a powerful approach to predict evolutionary outcomes when individuals interact, particularly in the presence of frequency-dependent selection, was widely adopted by behavioral ecologists (Krebs and Davies 1978) and continues to be influential in behavioral research (McNamara and Leimar 2020).

In the 1980s and 1990s, a body of theory incorporating explicit genetic considerations into social evolution models began to be developed. Lande (1980, 1981) and Kirkpatrick (1982) both formalized Fisher’s model of sexual selection and showed that the genetic correlations that arise between sexually selected traits and mating preferences can sometimes drive runaway evolution. Cheverud (1984), Lynch (1987), and Kirkpatrick and Lande (1989) incorporated maternal effects, which had long been studied by quantitative geneticists (Dickerson 1947; Willham 1963, 1972; Falconer 1965), into general evolutionary models. Echoing earlier work by Griffing (1967, 1977, 1981), Moore et al. (1997) extended maternal effects models to accommodate generalized social interactions between individuals, including unrelated individuals. Such “interacting phenotype” models depend on “indirect genetic effects” (IGEs), the modification of an individual’s phenotypic expression in response to genetically variable traits in a social interactant. IGEs, which are analogous to genetic maternal effects but which arise between any interacting conspecifics, may drastically alter responses to selection (Moore et al. 1997; Wolf et al. 1998; McGlothlin et al. 2010).

Another key aspect of quantitative genetic models of social evolution is social selection, which is caused when one individual’s traits affect the fitness of another and is thus also amenable to quantitative genetic modeling (Wolf et al. 1999). The term “social selection” has been used in different ways by many authors (West-Eberhard 1979, 1983, 2014; Roughgarden et al. 2006; Roughgarden 2012; Lyon and Montgomerie 2012), focusing on different behavioral mechanisms and contexts (e.g., competition for social partners versus negotiation of social dynamics). Common to all conceptions of social selection is that the fitness of a focal individual is a function of its social environment, which includes the traits of its partners. In quantitative genetics, social selection can be modeled as a selection gradient measuring the fitness effect of social traits on the fitness of a focal individual (Wolf et al. 1999), which is equivalent to neighbor-modulated or group-level selection (Queller 1992; Bijma et al. 2007). Regardless of how it is represented, social selection may lead to evolutionary change when related individuals interact or in the presence of IGEs (Bijma and Wade 2008; McGlothlin et al. 2010; Akçay and Van Cleve 2012).

Despite the long tradition of studying social evolution via evolutionary game theory and quantitative genetics, the two frameworks have remained mostly separate, at least in practice. This divide has persisted in part because the goals of the two approaches differ slightly. While game theory models usually focus on solving for an equilibrium ESS (Maynard Smith 1982; McNamara and Leimar 2020), the goal of a quantitative genetic model of social evolution is usually to develop predictions for short-term evolutionary response to selection (Lande 1980; Moore et al. 1997). Yet this apparent difference masks a deep complementarity. Solving for an ESS requires writing equations for short-term evolutionary change and calculating what phenotypes can resist it. Quantitative genetics is typically concerned with changes in trait values without regard to whether such change is near or far from an evolutionary equilibrium, but can also be used to predict which traits will be stable in the face of such change. Conversely, evolutionary game theory places emphasis on modeling the behavioral processes that result in differential fitness outcomes, while quantitative genetics remains largely agnostic about such processes. Thus, although evolutionary game theory and quantitative genetics usually differ in emphasis, they fundamentally describe the same processes. A synthesis of the two frameworks can provide general insight into social evolution by combining models that focus on predicting equilibria with those that describe the process required to reach such equilibria (Moore and Boake 1994).

Here we demonstrate parallels between evolutionary game theory and quantitative genetics that are not widely appreciated in social evolution. Our treatment complements earlier syntheses (Aoki 1983, 1984; Gomulkiewicz 1998; Lion 2018; Lehtonen 2018; Queller 1992, 2011; Abrams et al. 1993; Taylor 1996), but differs in that we aim to explicitly incorporate the parameters of ESS models into interacting phenotype theory and vice versa. First, we express fitness effects in a general evolutionary game in terms of quantitative genetic selection gradients, showing how payoff matrices relate to parameters that may be estimated in natural populations. Next, we consider how both short-term response to selection and long-term evolutionary outcomes may be affected by the presence of indirect genetic effects. We discuss two specific games, the prisoner’s dilemma and the hawk-dove game, as examples. Finally, we discuss the relevance of the IGE parameter *ψ*, which measures the strength and direction of phenotypic social interactions, for evolutionary games that include reciprocity (Trivers 1971; Axelrod and Hamilton 1981).

### Game Theory and Social Selection

In this section, we connect fitness effects in evolutionary game theory to the two selection gradients in interacting phenotype theory, the nonsocial (or direct) selection gradient *β*_*N*_, which measures the effects of an individual’s traits on its own fitness, and the social (or indirect) selection gradient *β*_*S*_, which measures the effects of the traits of others on an individual’s fitness (Wolf et al. 1999). Throughout, we assume that individuals interact in dyads, but our results can easily be extended to larger groups. Our treatment builds upon previous work by Taylor (1996), Taylor and Frank (1996), Queller (2011), Van Cleve (2017), Araya-Ajoy et al. (2020), and others, but we provide additional insights into the estimation of parameters in wild populations.

In game theory, the fitness costs and benefits of an evolutionary game are described by the payoff matrix, which defines the fitness consequences of an individual employing (or “playing”) a particular strategy in a given social context. The values in the payoff matrix can be written in the form *V* (*z*|*z*′), which represents the expected payoff of an individual playing strategy *z* against an individual (or group) playing strategy *z*′. In a simple two-player game where an individual may take one of two strategies, we can define *z* as a binary trait where one strategy is *z* = 1, which corresponds to a player using strategy “1” with probability one, and the second strategy is *z* = 0, which corresponds to a player using strategy “0” with probability one and strategy “1” with probability zero. A general payoff matrix that can accommodate a wide variety of two-player games can be written as

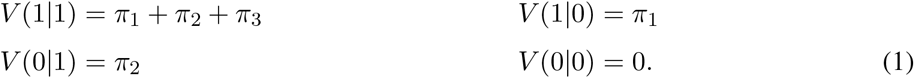

Here, the parameter *π*_1_ represents the additive payoff to the focal individual of playing strategy *z* = 1 and *π*_2_ represents the additive payoff of facing strategy *z*′ = 1. Parameter *π*_3_ represents nonlinearity or synergy, i.e., the additional payoff that accrues only when both players play strategy “1.” The payoffs in Equation 1 are assumed to be normalized with respect to the payoff *V* (0|0), which is set to zero. This decision is arbitrary; fitness payoffs may be written by normalizing with respect to any of the other cells of the matrix. In this way, this notation can encompass any two-player, two-strategy game (Queller 2011; Van Cleve 2017). The payoffs in Equation 1 can be used to write a general equation for the absolute fitness of an individual:

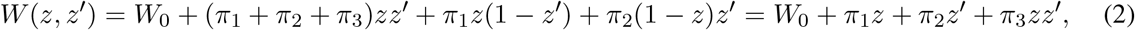

where *W*_0_ represents baseline fitness outside the context of the game. Using this fitness function, we can interpret *z* as before, a binary trait with values zero and one, or allow *z* to be a continuous trait with values within the interval [0, 1] where *z* is the probability of using strategy “1” and 1 − *z* is the probability of using strategy “0”. Further, we can use the fitness function *W* (*z, z*′) to calculate all the possible ESSs of the game using standard methods (Maynard Smith 1982; Hofbauer and Sigmund 1998).

Like evolutionary game theory, interacting phenotype theory also calculates fitness in a social context but using the notation and tools of quantitative genetics. Specifically, the interacting phenotype theory fitness function allows one to partition total selection into individual and socially mediated components (Wolf et al. 1999). Quantitative genetic models often start with relative fitness, which is defined as absolute fitness divided by mean fitness, or 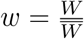. For a one-trait model, relative fitness is modeled as

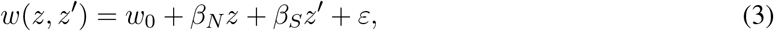

where *w*_0_ is an intercept, *z* represents the phenotype of the focal individual, *z*′ represents the phenotype of its social partner, and *ε* is a normally-distributed error term with an expectation of zero. The effect of the focal phenotype on fitness is by the nonsocial selection gradient (*β*_*N*_), and the effect of the social phenotype is captured by the social selection (*β*_*S*_). These selection gradients are partial regression coefficients and represent partial fitness effects, i.e., the effect of a given phenotype on fitness while holding the other constant. When group means are used in place of *z*′, this partitioning is identical to the approach known as contextual analysis (Goodnight et al. 1992; Heisler and Damuth 1987), and when that group mean excludes the focal individual, it is identical to the neighbor model of group selection (Lehtonen 2020; Okasha 2004, 2006). Nonsocial and social selection gradients can be estimated using a simple modification of the multiple regression method of Lande and Arnold (1983) and thus provide a useful tool for studying social evolution in natural populations (Wolf et al. 1999; Formica et al. 2011).

Fitness is the common currency of the two bodies of theory and so provides a translation between them. We first assume that the payoff matrix (Equation 1), which we have defined in terms of discrete binary strategies, can be used to define a continuous surface of fitness effects defined by continuous traits *z* and *z*′. These could represent the “mixed strategies” of game theory, where *z* represents the probability of performing strategy 1. In this case, the payoff matrix (Equation 1) would represent the four corners of a bounded fitness surface. However, *z* and *z*′ could also represent other continuous traits that take values beyond the interval [0, 1]. In the latter case, the four points in the payoff matrix now define a fitness function that may be used to calculate the fitness of individuals with any trait value.

The selection gradients in Equation 3 can now be calculated by differentiating Equation 2 with respect to focal and social phenotypes. Following Taylor and Frank (1996, see also Lande and Arnold 1983; Charlesworth 1990; Iwasa et al. 1991; Taper and Case 1992; Abrams et al. 1993; Taylor 1996), the regression coefficients can be expressed as

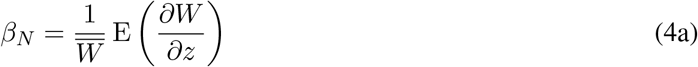

and

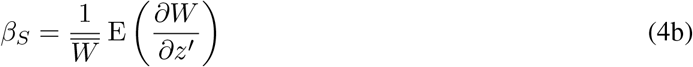

when we assume that *z* and *z*′ are jointly normally distributed. Equations 4 represent expectations of partial derivatives of the absolute fitness function divided by population mean fitness. These expectations may be approximated by evaluating derivatives at the population mean, or

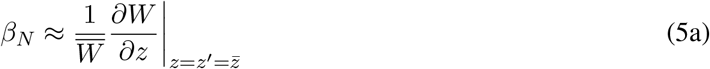

and

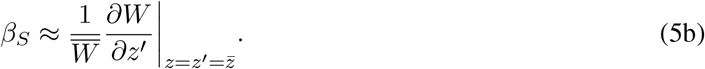

These expressions are exact when *W* (*z, z*′) is at most quadratic in either *z* or *z*′, as is the case in Equation 2, and they hold more generally whenever the phenotypic variance is small (Abrams et al. 1993) or when selection is weak (Taylor 1996).

Using Equations 2 and 5, we can calculate the nonsocial and social selection gradients in the two-player game defined by Equation 1 as

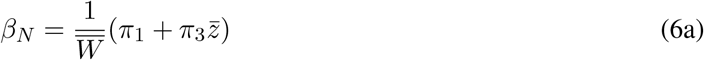

and

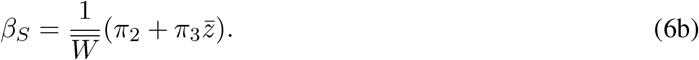

where the average fitness is

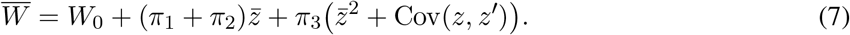

The covariance term in Equation 7, which will be nonzero when individuals are related, when individuals assort on the basis of phenotype, or when there are indirect genetic effects (i.e., when one individual’s phenotype is adjusted in response to its social partner), is equal to the interactant covariance (*C*^*ij*^′) of Wolf et al. (1999).

Equations (6-7) show that the nonsocial and social selection gradients from interacting phenotype theory are not necessarily constant across generations but rather should change as the population mean changes. In the absence of nonlinear payoffs (*π*_3_ = 0), these changes will derive solely from changes in mean fitness (equation 7), which will not influence the signs of the selection gradients and will be small when selection is weak. However, in any evolutionary game with nonlinear payoffs (*π*_3_ ≠ 0), both nonsocial and social gradients may change in sign as well as magnitude (Figure 1).

**Figure 1:**
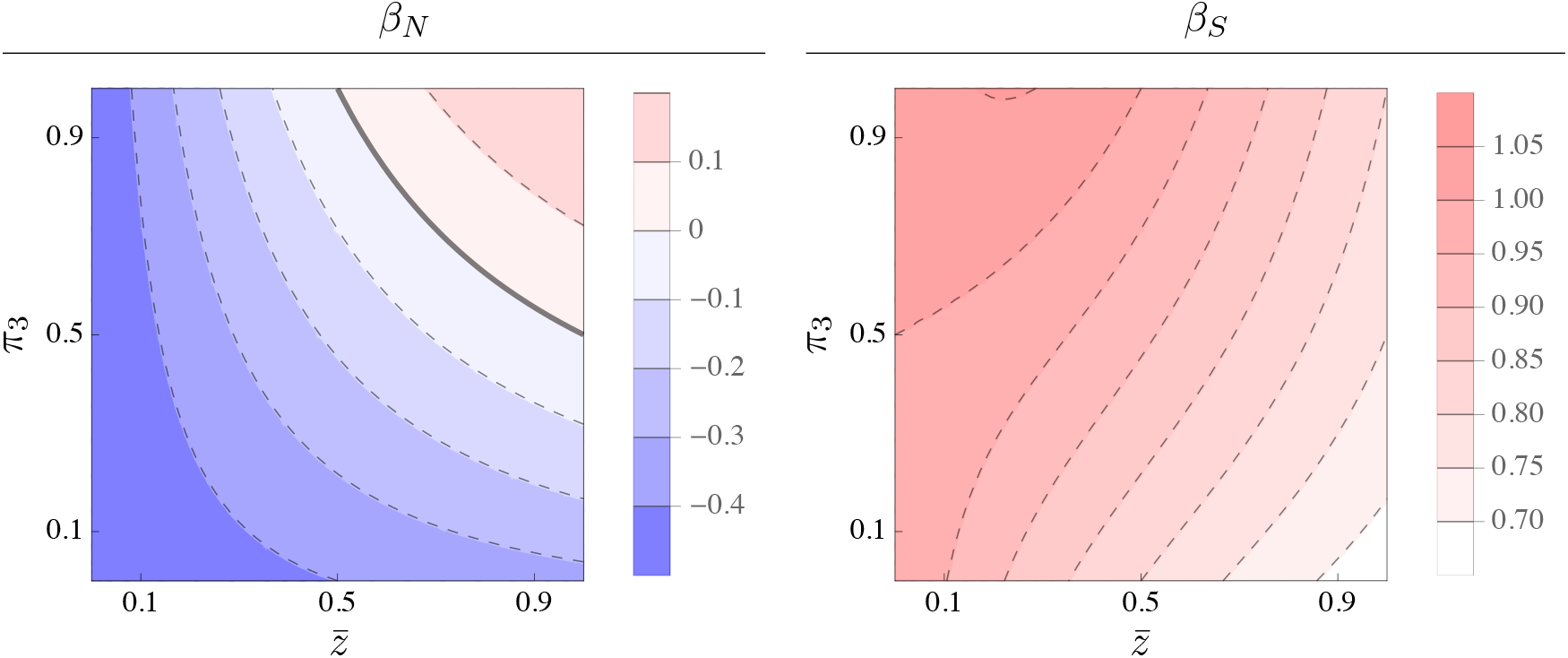
Dependence of nonsocial (*β*_*N*_) and social (*β*_*S*_) selection gradients on the trait mean 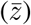 and the nonlinear payoff *π*_3_. Trait *z* is costly to self (*π*_1_ = −0.5) but beneficial to others (*π*_2_ = 1.0). Average fitness is calculated using Equation 7 where *W*_0_ = 1 and the covariance term is set to zero. When synergistic effects are small (*π*_3_ *< π*_1_), the selection gradients change in magnitude with 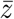 but do not change in sign. When fitness effects are synergistic enough (*π*_3_ *> π*_1_), however, the nonsocial selection changes in sign once 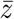 is large enough. This effect results in the trait changing from representing a net cost to self to a net benefit to self.

When nonlinear fitness effects are absent or weak (i.e., when |*π*_3_| *< π*_1_ and |*π*_3_| *< π*_2_), the interpretation of nonsocial and social selection gradients does not differ from the interpretation of the additive terms in the payoff matrix. For example, a trait that represents a cost to self (*π*_1_ *<* 0 in the payoff matrix will always result in a negative nonsocial selection gradient (Figure 1). However, when fitness effects are strongly synergistic (i.e., when |*π*_3_| *> π*_1_ or |*π*_3_| *> π*_2_), our interpretations of the two types of parameters may diverge. A strongly positive *π*_3_ coupled with a negative *π*_1_, for example, may result in a nonsocial selection gradient that is negative when the trait mean is low and positive when the trait mean is high (Figure 1). The interpretation here depends on perspective. A game theorist might view this trait as lacking “strategic dominance” (Binmore 2007) since the cost of switching strategies, *V* (1|1)−*V* (0|1) = *π*_1_+*π*_3_ and *V* (1|0)− *V* (0|0) = *π*_1_, have different signs, whereas a quantitative geneticist would note that the trait changes from representing a net cost to a net benefit depending on the trait mean.

Our results show that interacting phenotypes models potentially include two types of frequency dependence. The first occurs whenever there is social selection, i.e., whenever *π*_2_ and *β*_*S*_ are nonzero. In this case, the fitness of a given phenotype depends upon the individual (or individuals in a group model) with which it interacts (Wolf et al. 1999). This contribution can be called “global frequency dependence” because it creates variance in fitness across pairs (or groups) of interactants existing in the same population at the same time. Local frequency dependence aligns with the concept of frequency dependence in game theory and population genetics, which is usually defined to include any type of socially dependent fitness effect (McNamara and Leimar 2020). Local frequency dependence should be present in any evolutionary game or social selection model. The second type occurs when the sign of selection gradients changes across generations with the trait mean as in Figure 1. This contribution can be called “global frequency dependence” because the total effect of selection on the trait, 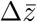, depends upon the makeup of the whole population. Global frequency dependence aligns with the definition of frequency-dependent selection in quantitative genetics, i.e., a change in the direction of selection with a change in the phenotypic mean (Lande 1976). Global frequency dependence will arise only when nonlinear payoffs (*π*_3_) exist.

Previous interacting phenotype models have obscured the contribution of nonlinear fitness effects, which has been cited as one of their primary limitations (Westneat 2012; Araya-Ajoy et al. 2020). As noted by Araya-Ajoy et al. (2020), one of the most important consequences of nonlinear fitness effects is that they can lead to evolutionary feedback. In other words, because the strength of selection may change with the trait mean in the presence of synergy, the rate of evolution of a social phenotype may potentially increase each generation. Although the inclusion of nonlinear effects does not change the predictions of short-term response to selection (Moore et al. 1997; Bijma and Wade 2008; McGlothlin et al. 2010), it is important to consider how nonlinear effects might change selection gradients when attempting to draw conclusions about social evolution over many generations.

An alternative approach (Queller 1985, 2011; Westneat 2012; Araya-Ajoy et al. 2020) partitions selection to include a separate gradient (*β*_*I*_) that estimates the interactive (i.e., multiplicative) effect of focal and social traits (*zz*′). In this partitioning, the three selection gradients take on expectations that are independent of the mean, with 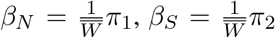, and 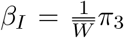. However, we prefer the standard definitions of *β*_*N*_ and *β*_*S*_ in Equation 6. As we will show in the next section, using standard definitions allows the incorporation of fitness effects from any evolutionary game directly into equations for evolutionary response (Bijma and Wade 2008; McGlothlin et al. 2010). In addition, these definitions directly correspond to the selection gradients that would be measured in a social selection analysis of a natural population (Wolf et al. 1999).

With a sufficiently large dataset one could estimate nonlinear fitness effects in a natural population by expanding the Wolf et al. (1999) model to include quadratic terms (Lande and Arnold 1983; Phillips and Arnold 1989). Defining relative fitness as

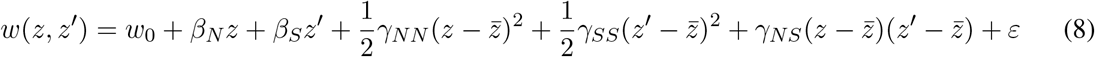

has the advantage of estimating evolutionarily relevant selection gradients while estimating the total curvature of the fitness surface (*γ*_*ij*_ terms). Here, the nonsocial-social correlational selection gradient (*γ*_*NS*_) captures the same fitness interaction as *β*_*I*_ (Westneat 2012; Araya-Ajoy et al. 2020), while the directional gradients include all fitness effects relevant to evolutionary response.

### Response to Selection in Evolutionary Games

One advantage of defining selection gradients in terms of the payoff matrix is that it enables calculation of the predicted response to selection that arises from an evolutionary game. In general, the predicted response to selection for a socially influenced trait can be calculated from the Price (1970, 1972) equation using Equation 3 as

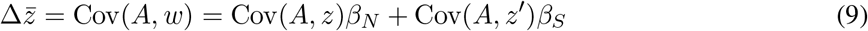

(Frank 1997; McGlothlin et al. 2010), where *A* is the total breeding value of trait *z* in an individual. This equation is useful because it partitions the relationship between breeding values and phenotypes (the two covariances) from the relationship between phenotypes and fitness (the two selection gradients). The two covariance terms on the right-hand side of Equation 9 represent two types of genotype-phenotype relationships. The first describes the relationship between an individual’s own genotype and phenotype, and the second describes whether an individual’s genotype is associated with phenotypes found in its social environment.

To calculated these covariances, we must specify a genotype-phenotype model. One common model for the genetics of interacting phenotypes in quantitative genetics partitions a single phenotype *z* into the influence deriving from a focal individual’s own additive genetic (*a*) and environmental (*e*) contributions and the effect of a social partner’s phenotype (*z*′):

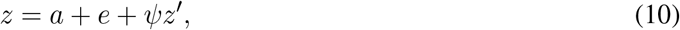

where *ψ*, which is a constant representing the importance of the social interaction for the expressed trait, theoretically ranges from -1 to 1 for a single-trait model (Moore et al. 1997). The coefficient *ψ* is a way to incorporate phenotypic responsiveness to the social environment into quantitative genetic models. Specifically, when *ψ* is nonzero, the expression of phenotype *z* depends in part on a response to the trait of its social partner (Moore et al. 1997). This can arise, for example, if the phenotype is behavioral and individuals observe and respond to each other’s behavior in real time; *ψ* then represents this behavioral responsiveness (Akçay et al. 2009; Akçay and Van Cleve 2012). The model in Equation 10 may be used to develop expressions for both an individual’s breeding value 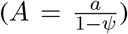 and the covariances in Equation 9 (Moore et al. 1997; McGlothlin et al. *2010*). *Whenever z* is heritable and *ψ* ≠ 0, the total covariance between breeding values and phenotype will depend upon two sources: direct genetic effects caused by an individual’s own genes and IGEs caused by the social partner’s genes and mediated by *ψ*.

Because the social partner’s trait is also influenced by genetic, environmental, and social components, Equation 10 can be expanded as

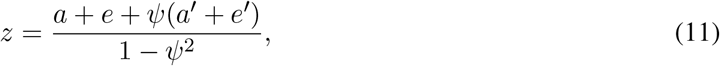

where the denominator now represents the degree of feedback between the two interactants (Moore et al. 1997). Equation 11 shows that phenotypic expression involves two genetic effects: a direct genetic effect proportional to *a* and an indirect genetic effect (IGE) proportional to *a*′. If relatives interact or pairs form nonrandomly, these effects may be correlated. We model the correlation between the breeding values of different individuals using a genetic relatedness coefficient *r*. The predicted response to selection in the presence of both relatedness and IGEs can be shown to be

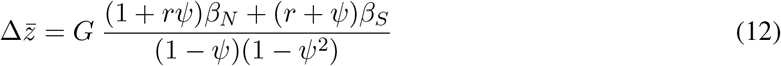

(McGlothlin et al. 2010; cf. Akçay and Van Cleve 2012), where *G* represents additive genetic variance.

An equally valid alternative to this trait-based model partitions the total breeding value (*A*) of an individual trait into direct and indirect components without reference to a specific phenotype of the social partner and uses variance components to predict the response to selection (Griffing 1967; Bijma et al. 2007; Bijma and Wade 2008; McGlothlin and Brodie 2009; Bijma 2014). While trait-based IGE models, such as the one we develop here, decompose IGEs into effects of specific traits, variance-component models include terms that encompass all IGEs on a particular trait. We use a univariate trait-based model here to parallel game theory approaches, but this is easily generalizable to a multivariate scenario. Including effects of other traits on evolutionary response requires either a variance-component model (Bijma and Wade 2008) or a multivariate trait-based model (Moore et al. 1997).

Equation 12, when combined with our previous definitions of nonsocial and social selection gradients (Equation 6), allows the prediction of short-term evolutionary change from a wide variety of evolutionary games. In addition, the incorporation of both relatedness and phenotypic responsiveness into Equation 12 allows us to determine when these phenomena may affect both short-term evolutionary response and predicted evolutionary outcomes. In particular, *ψ* may be useful for modeling any game where strategies are allowed to change in response to the strategy of the opponent. Such phenotypic responsiveness is known variously as “(direct) reciprocity” (Trivers 1971; Axelrod and Hamilton 1981; Alexander 1985) or “negotiation” (McNamara et al. 1999). An alternative approach to the one we take here is to explicitly model how responsiveness affects the trait value over repeated rounds of interaction and incorporate those effects into the payoff matrix (Van Cleve and Akçay 2014; Van Cleve 2017). The variable used by Van Cleve and Akçay (2014) and Van Cleve (2017) to quantify responsiveness (*ρ*) is conceptually similar but not identical to *ψ* (Akçay and Van Cleve 2012). An advantage of our current approach is that it partitions the effect of selection on trait values from genetics, which is consistent with standard quantitative genetic theory and thus allows for the estimation of relevant parameters using quantitative genetic methods.

Equations (9-12) can also be used to predict the direction of evolution and evolutionary equilibria for a given model by setting 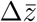 equal to zero and rearranging. For example, in the presence of both relatedness and IGEs, 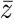 should evolve in a positive direction when

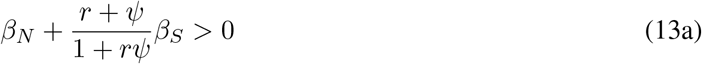

and should be at equilibrium when

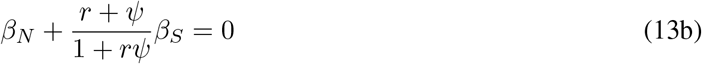

(McGlothlin et al. 2010, 2014; Akçay and Van Cleve 2012). In most cases, this approach should predict the ESS for a given evolutionary game (Taylor and Frank 1996). To explicitly illustrate how game theory and interacting phenotypes models can apply to the same behavioral scenarios, we analyze two simple evolutionary games using this approach below.

#### Example 1: Prisoner’s Dilemma

One classic evolutionary game is the prisoner’s dilemma (Rapoport and Chammah 1965), which is defined by the matrix

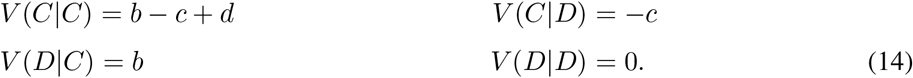

In this game, each player may either cooperate (*C*) with the other player, in which case it pays a fitness cost *c*, or defect (*D*), acting in its own interest and paying no fitness cost. Interacting with a cooperator leads to a fitness benefit *b*. The synergy term *d* is greater than zero when simultaneous cooperation yields additional fitness benefits. Most evolutionary games can be recovered from the prisoner’s dilemma by varying the signs and relative magnitudes of *b, c*, and *d* (Van Cleve and Akçay 2014; Van Cleve 2017). Indeed, the payoff matrix represented in Equation 14 is identical to that in Equation 1 except for signs.

Using Equations 5, the nonsocial and social selection gradients in this game are

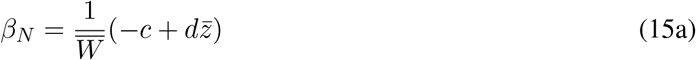

and

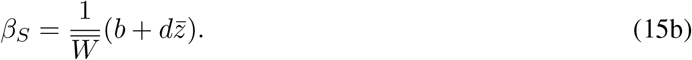

As expected, nonsocial and social selection are proportional to the costs and benefits of cooperation in the absence of synergy (*d* = 0) and depend on the population mean in the presence of synergy (*d* ≠ 0). In the absence of IGEs and relatedness, evolution depends only upon nonsocial selection, and cooperation will always be disfavored in the absence of positive synergy. Even with positive synergy, however, cooperation is unable to increase when 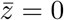.

Adding IGEs and relatedness allows us to derive a version of Hamilton’s rule that includes synergy. Cooperation will increase (i.e., 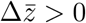) when

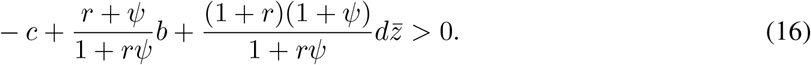

As in previous versions of Hamilton’s rule that did not explicitly incorporate synergy (McGlothlin et al. 2010, 2014), Equation 16 shows that relatedness and IGEs have completely symmetrical effects on the evolution of cooperation (Akçay and Van Cleve 2012; Van Cleve and Akçay 2014; Van Cleve 2017). The influence of fitness synergy on the evolution of altruism has been discussed at length elsewhere (e.g. Queller 1985). Two additional insights arise from including IGEs into this model, however. First, when *ψ >* 0, IGEs may facilitate the invasion of cooperation when 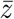 is close to zero by creating a phenotypic covariance between interactants and increasing the effect of the *b* term (i.e., the linear component of social selection). Second, IGEs may act in concert with relatedness to enhance the effects of positive synergy as cooperation and 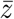 increase.

#### Example 2: The Hawk-Dove Game

We next consider the classic hawk-dove game, which involves two conspecifics competing over a resource (Maynard Smith and Price 1973; Maynard Smith 1982). When two doves meet, they divide the resource or decide the contest without aggression. When two hawks meet, they fight to determine the contest, with one hawk winning and the other paying a cost. When a hawk meets a dove, the dove flees and the hawk takes the resource. In this game, the payoffs can be expressed as

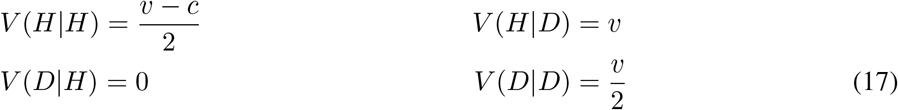

where *H* represents the hawk strategy, *D* represents the dove strategy, *v* represents the value of the resource, and *c* represents the cost of fighting over that resource. Fitness can be expressed as a function by equating *z* = 1 with a pure hawk strategy and *z* = 0 with pure dove:

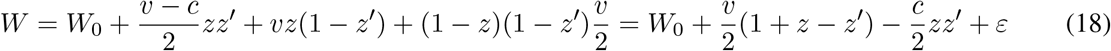

As before, we can translate this fitness function into selection gradients by taking partial derivatives of Equation 18, yielding

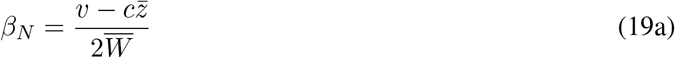

and

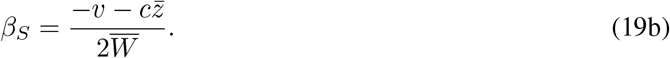

To predict evolutionary rates and solve for equilibria, we substitute Equations 19 into Equation 12. In the absence of IGEs and relatedness, the result reduces to

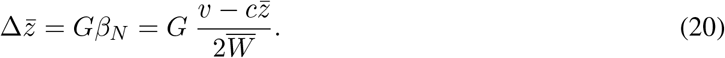

Solving for equilibrium,

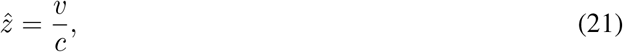

which corresponds to the classic ESS for the continuous hawk-dove game (Maynard Smith 1982). Naturally this mixed-strategy equilibrium is stable only when *v < c* (so that 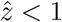). When 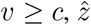 from Equation 21 is unstable, and 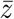 will evolve to its boundary condition at 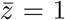, which is the ESS in the hawk-dove game when hawk is the dominant strategy.

Now imagine that the two individuals are allowed to adjust their behavior in response each other. Such negotiation can be modeled as an IGE. By substituting Equations 19 into Equation 12 and setting only relatedness to zero, we find a predicted response to selection of

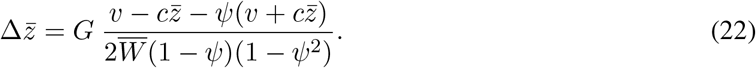

We can solve for the equilibrium phenotype by setting the left-hand side equal to zero and solving for 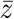:

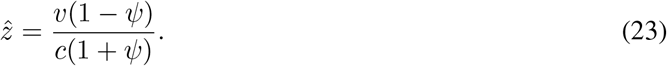

Equation 23 shows that phenotypic adjustment changes the predicted equilibrium of a hawk-dove game. If *ψ >* 0, there is positive feedback, meaning that interactants will adjust their behavior based on what the other does, which results in lower equilibrium hawkishness than predicted in the phenotypic model. This result is equivalent to Grafen’s (1979) result for relatedness, with *ψ* in place of *r*. In contrast, if *ψ <* 0, individuals become more hawkish when their opponent is more dovish, and vice versa. This results in a higher equilibrium hawkishness.

### Indirect Genetic Effects and Reciprocity

Above, we have argued that IGEs may be used as a general way to model phenotypic adjustment in evolutionary games. Perhaps the most well-known type of phenotypic adjustment is reciprocity or reciprocal altruism, first discussed by Trivers (1971). A typical model of reciprocity involves sequential interactions between individuals where memory of one or more previous interactions determines the action taken in the current interaction. It has been argued that IGEs allow quantitative genetic models to incorporate reciprocity (Bleakley and Brodie 2009; McGlothlin et al. 2010), but this relationship has not been made mathematically.

In evolutionary game theory, reciprocity is often modeled using a repeated or iterated version of the prisoner’s dilemma game (Axelrod and Hamilton 1981; McElreath and Boyd 2007). After meeting once and playing a prisoner’s dilemma, two individuals play again with some probability. By allowing various repeated strategies to compete in a computer simulation, Axelrod and Hamilton (1981) showed that a strong strategy in this game is tit-for-tat, where each player mimics its partner’s behavior from the previous round. A population that employs a tit-for-tat strategy can prevent the invasion of cheaters, because an individual that cheats on the first meeting is punished in later iterations.

This game appears to match a typical IGE model with one exception. The evolutionary game theory model assumes that interactions are repeated and sequential, while IGE models involve a single interaction with phenotypic feedback. To show the relationship between the two models, we can express the phenotypes for a pair of individuals that interact sequentially using a quantitative genetic model in which subsequent interactions are included as different traits. Suppose that in the first interaction, two individuals express a phenotype that is unaffected by the other individual:

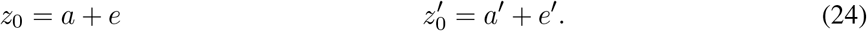

In Equations 24 and those that follow, the subscript refers to the number of times two individuals have previously met (i.e., *z*_0_ and 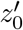 are phenotypes for two individuals that have not previously interacted). When two individuals interact again, each individual adjusts its phenotype based on what occurred in their last interaction. We use *y* to represent the fraction of the phenotype that is attributable to this adjustment, while 1−*y* is the fraction of the phenotype based on the individual’s own genes and environment. A value of *y* = 1 would be equivalent to a perfect tit-for-tat response. For simplicity, we assume that *y* is a population parameter. However, the model could easily be expanded to allow each individual to have its own value of *y*. Thus, at the second interaction, the two phenotypes can be written as

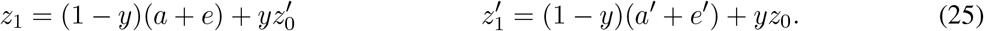

Because all subsequent interactions follow the same form, this can be generalized as

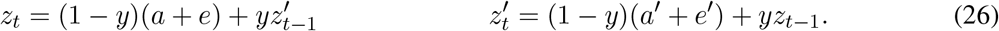

Now suppose that each additional interaction occurs with probability *p*. To make a direct comparison to the standard IGE model, we measure the phenotype as the average behavior across all repeated interactions between the two individuals. The average phenotype is the sum of values across all steps divided by the total number of interactions, (1 − *p*)^−1^, or

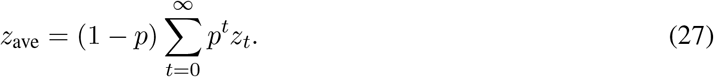

In the Appendix, we show that Equation 27 can be expressed as

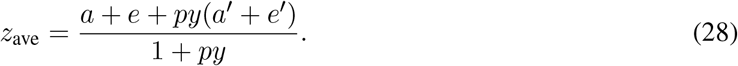

The numerator in Equation 28 corresponds to the phenotypic definition from the standard IGE model (Equation 11), with the equivalency

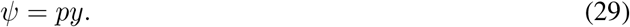

In other words, the the interaction effect *ψ* is equal to the probability of repeated interactions multiplied by the strength of reciprocal response. However, the sequential model differs slightly from the standard IGE model; namely, there is an additional factor of (1 − *ψ*) in the standard model. In Equation 28, as *py* approaches 1, the denominator approaches 2 and the numerator approaches the sum of the initial values of the two interactants. This shows that stronger, more likely interactions cause the phenotype to resemble the average of the initial values of the two interactants.

To derive an equation for the response to selection in this model, we take the population average of Equation 28,

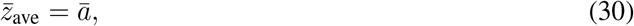

which shows that unlike in the standard IGE model, the population mean equals the mean additive genetic value. This allows us to solve for a response to selection by using the additive genetic value as the breeding value in Equation 9, yielding

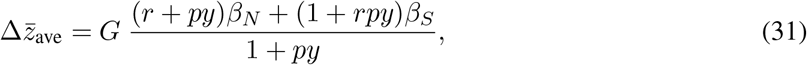

which, with *ψ* = *py*, differs from Equation 12 by a factor of (1 − *py*)^−2^.

Equation 31 shows that the predicted response to selection when repeated interactions occur will be proportional to the predicted response for a single interaction with simultaneous feedback; however, assuming *py >* 0, the rate of evolutionary change will be reduced. Thus, when we assume that interactions occur sequentially instead of simultaneously, likely a more realistic assumption, the runaway feedback effect that normally characterizes models with reciprocal IGEs (Moore et al. 1997; McGlothlin et al. 2010) disappears. The response to selection still depends on both direct genetic effects and IGEs in a similar way, but the rate of evolution no longer spirals to infinity as the strength of the IGE increases.

Despite the difference in predicted evolutionary rate between the two models, other conclusions depend only on the numerator and will not differ. This allows us to use estimates of *ψ* to make general predictions about the direction of evolution no matter what model the interactions follow. For example, the condition for the evolution of reciprocity with sequential interactions can be given as a form of Hamilton’s rule:

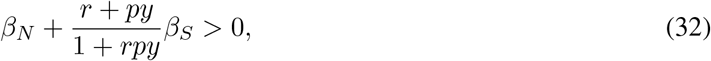

which with *ψ* = *py* is identical to Equation 13a. When non-relatives interact, Equation 32 simplifies to

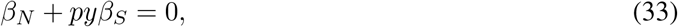

which reproduces results from prior models of reciprocity and responsiveness (André and Day 2007; Akçay et al. 2009). Equations 32-33 indicate that, as expected, the evolution of reciprocity depends on *py* or approximately, *ψ*. The evolution of altruistic behavior is favored when repeated encounters with the same individual are more likely (higher *p*) and when the response more closely resembles tit-for-tat (higher *y*). This result corresponds to the standard result that the evolution of reciprocity via a tit-for-tat strategy is favored when subsequent encounters are more likely (McElreath and Boyd 2007). It is important to note, however, that our model assumes that the parameters *p* and *y*, and consequently, *ψ*, are fixed, and conclusions may differ when *ψ* is allowed to evolve (André and Day 2007; Akçay et al. 2009; Kazancioglu et al. 2012; André 2015). The assumption of fixed *p* and *y*, or *ψ*, allows prediction of short-term evolution.

## Discussion

Our analysis highlights important connections between evolutionary game theory and quantitative genetic models of interacting phenotypes. Specifically, the fitness effects that arise from the payoff matrix in social games translate directly into the nonsocial and social selection gradients of interacting phenotypes theory from quantitative genetics. Various types of phenotypic modification can be modeled using the interaction coefficient *ψ*, which quantifies the strength of IGEs. These relationships make it straightforward to incorporate any type of interaction described by an evolutionary game into the predictive equations for evolutionary responses to selection. Thus, evolutionary models developed under the “phenotypic gambit” can be translated into explicitly genetic models simply by specifying the genotype-phenotype relationship typical of quantitative genetics, namely the partitioning of phenotype into direct and indirect genetic components and an environmental component.

The primary advantage of translating between game theory and quantitative genetics is that genetic models are typically formulated in terms of parameters that can be estimated in natural populations. For example, *β*_*N*_ and *β*_*S*_ can be estimated in natural populations using a modification of the standard regression method of Lande and Arnold (1983) that incorporates social phenotypes or group means as predictors (Heisler and Damuth 1987; Goodnight et al. 1992; Wolf et al. 1999). Studies that measure social selection have begun to accumulate (Formica et al. 2011; Farine and Sheldon 2015; Fisher and Pruitt 2019; Santostefano et al. 2020), and we hope this work encourages empiricists studying social phenotypes in natural populations, particularly those whose questions are motivated by evolutionary game theory, to estimate *β*_*S*_ for their traits of interest. Such estimates may be used to parameterize payoff matrices, providing strong tests of predictions from game theory. As we note above, correlational selection gradients may also be incorporated to test for the presence of synergistic interactions by using Equation 8.

We argue that the phenotypic modification captured in evolutionary games leads to IGEs, which may also be estimated in natural or laboratory populations. The parameters of variance-component models of IGEs, i.e., direct and indirect genetic (co)variance (Bijma et al. 2007; Bijma and Wade 2008), may be estimated using quantitative genetic animal models (Bijma 2010; Wilson et al. 2009, 2010, 2011). Methods also exist to estimate *ψ*, either by using inbred lines or test strains (Bleakley and Brodie 2009) or as functions of variance components (McGlothlin and Brodie 2009). Estimates of IGEs are increasingly common, particularly in populations of domestic animals (Wade et al. 2010), and a number of studies have estimated IGEs specifically for behavioral traits that have often been analyzed using game theory, such as aggression (Wilson et al. 2009, 2011; Saltz 2013; Alemu et al. 2014; Santostefano et al. 2017; Han et al. 2018; Lane et al. 2020). Despite this increased interest in IGEs, however, studies attempting to estimate *ψ* remain rare (but see Bleakley and Brodie 2009; Edenbrow et al. 2017). We hope that our results motivate empiricists to estimate this parameter more often, particularly when testing for the phenomenon of reciprocity.

Our results show that game theory holds lessons for quantitative geneticists as well. Nonsocial and social selection gradients represent net fitness effects of a phenotype. As such, they are often interpreted as quantitative genetic analogs of Hamilton’s (1964a) costs and benefits. This equivalence may be accurate in some cases, but there are several scenarios in which the two diverge (McGlothlin et al. 2014; Hadfield and Thomson 2017). As our results show, one such case arises when strong nonlinear fitness effects exist (see also Westneat 2012; Araya-Ajoy et al. 2020). We urge empiricists to exercise caution when mapping selection gradients to cost and benefit terms from evolutionary games, particularly when the correlational selection gradient *γ*_*NS*_ is nonzero. We note that selection gradients can always be correctly interpreted as the effective relative fitness cost or benefit of a measured phenotype in the current population, and they can be used to accurately predict short-term evolutionary response of socially influenced traits (Bijma and Wade 2008; McGlothlin et al. 2010) even when nonlinear effects are present.

The interaction coefficient *ψ* should also be interpreted with caution, particularly when predicting the rate of evolutionary change. Explicitly modeling phenotypic modification with sequential social interactions similar to prior models of direct reciprocity in the prisoner’s dilemma shows that the runaway feedback effects predicted by previous IGE models (Moore et al. 1997; McGlothlin et al. 2010) often may be attenuated. However, our treatment shows that other conclusions of the general model hold true because the direction of evolutionary response does not change. In particular, the sequential interaction model leads to a version of Hamilton’s rule identical to that of the standard model, indicating that *ψ* can be used to predict the evolution of reciprocity. In addition, methods to estimate *ψ* as a function of variance components are unaffected by the specific model of social interaction because feedback effects influence all variance components equally and thus cancel out (McGlothlin and Brodie 2009). Therefore, when it may be measured, *ψ* remains a useful estimate for the strength of reciprocity. Finally, *ψ* is likely to evolve (Chenoweth et al. 2010), which may also alter long-term evolutionary predictions (Akçay and Van Cleve 2012; Kazancioglu et al. 2012). A future contribution will address the conditions under which *ψ* will change in response to selection.

In conclusion, our results emphasize the benefits of unifying models of evolutionary outcomes and evolutionary process. Incorporating both types of theory leads to a holistic view of social evolution on microevolutionary and macroevolutionary scales. Further, both types of theory can contribute in different ways to the crucial feedback between theory and experiment. As we have argued, empirical quantitative genetics can estimate parameters to test and refine game theory models. At the same time, game theory can both provide conceptual context for the interpretation of theoretical and empirical quantitative genetic results and generate predictions about potential evolutionary outcomes that inform the design of microevolutionary studies of natural populations. We hope that our results will inspire further integration of game theory and quantitative genetics to lead to a richer understanding of the evolution of social phenotypes.

## Acknowledgments

Many thanks to Kim Hughes and Anjanette Baker for their efforts organizing the 2020 AGA Presidential Symposium. We thank Yimen Araya-Ajoy and David Westneat for helpful conversations and two anonymous reviewers for comments on the manuscript. JVC acknowledges NSF CAREER Award DEB-1846260 for support.

## Appendix

Here we will show how to obtain Equation 28 from Equation 27. First, we solve for the infinite sum, which we write as *z*_Σ_. Substituting for the phenotype at *t* = 0, when only the individual’s own genes and environment are important (Equation 24) yields

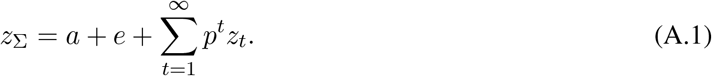

Starting with *t* = 1 (Equation 26), the phenotype depends on a combination of an individual’s own genes and environment (multiplied by the quantity 1 − *y*) and a response to its social partner’s previous phenotype (multiplied by *y*). Thus, the infinite series follows the form

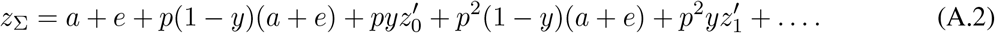

The infinite series in Equation A.2 is then

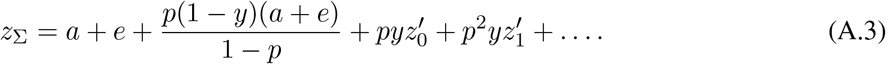

Next, we expand the contribution of the social partner, closing the sum in a similar way:

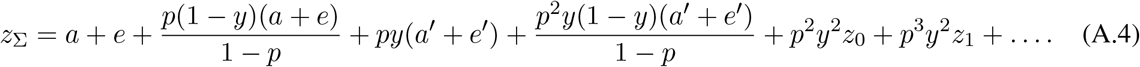

This leaves a portion of the phenotype due to feedback effects, which can be rewritten as

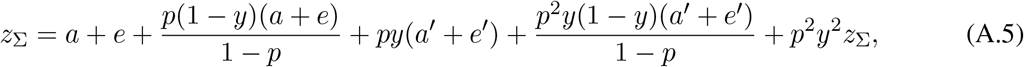

allowing us to close the entire sum as

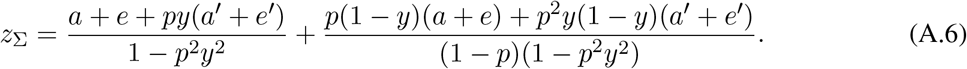

Simplifying,

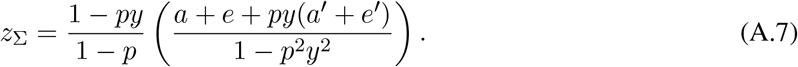

Now, we take the average across all interactions by substituting Equation A.7 into Equation 27, yielding

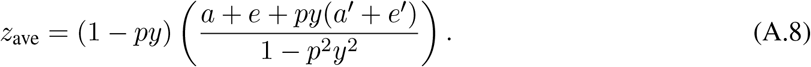

Finally, Equation A.8 easily simplifies to Equation 28.

